# Macroevolutionary Analyses Provide New Evidences of Phasmid Wings Evolution as a Reversible Process

**DOI:** 10.1101/2020.10.14.336354

**Authors:** Giobbe Forni, Jacopo Martelossi, Pablo Valero, Frank H. Hennemann, Oskar Conle, Andrea Luchetti, Barbara Mantovani

## Abstract

The concept that complex ancestral traits can never be re-acquired after their loss has grown popular since its initial formulation and it’s often referred to as Dollo’s law. Nonetheless, several macroevolutionary evidences - along with molecular ones - suggest instances where complex phenotypes could have been lost throughout a clade evolutionary history and subsequently reverted to their former state in derived lineages. One of the first and most notable rejection of Dollo’s law is represented by wing evolution in phasmids: this polyneopteran order of insects - which comprises stick and leaf insects - has played a central role in initiating a long-standing debate on the topic. In this study, a novel and comprehensive time-tree - including over 300 Phasmatodea species - is used as a framework for investigating wing’s evolutionary patterns in the clade. Despite accounting for several possible biases and sources of uncertainty, macroevolutionary analyses consistently support a dynamic and reversible evolution of wings, with multiple transitions to ancestral states taking place after their loss. Our findings suggest that wings and flight are decoupled from Phasmatodea diversification dynamics and that brachyptery is an unstable state, unless when co-opted for non-aerodynamic adaptations. We also explored how different assumptions of wings’ reversals probability could impact their inference: we found that until reversals are assumed to be over 30 times more unlikely than losses, they are consistently retrieved despite uncertainty in tree and model parameters. Our findings demonstrate that wings evolution can be a reversible and dynamic process in phasmids and contribute to shape our understanding of how complex phenotypes evolve.

## Introduction

Traits are commonly lost during evolution and - although some changes can be easily reverted in the short time (Teotónio and Rose 2000; Rebolleda and Travisano 2019) - it can be argued that the loss of complex ones is irreversible over long time spans. This concept is often referred to as Dollo’s law (Dollo 1893), despite in its original conception reversals of complex traits were considered possible as secondary convergence events (Gould 1970). Nonetheless, the concept that complex structures lost in the line of evolution cannot revert back to their former state later in the lineage has become popular, due to its intuitiveness and the frequent examples supporting it (Collin and Miglietta 2008). Although this evolutionary principle is still commonly accepted, an increasing number of cases where it is apparently violated have been proposed.

Most challenges to Dollo’s law come from macroevolutionary approaches and include a large number of examples, such as shell coiling in limpets (Collin and Cipriani 2003), compound eyes in ostracods (Syme and Oakley 2012), sex and parasitism in mites (Klimov and O’Connor 2013; Domes et al. 2007), mandibular teeth in frogs (Wiens 2011), limb evolution (Kohlsdorf and Wagner 2006), eggshells and oviparity (Lynch and Wagner 2010; Recknagel et al. 2018; Esquerré et al. 2020) in Squamata. At the same time, several possible mechansisms underlying the reversible evolution of complex traits, have been proposed, including phenotypic plasticity (Visser et al. 2021; Parker et al. 2021) and reticulate evolution (Horreo et al. 2020). Examples of compensatory mutations which have been able to rescue the once-lost functionality of genes have also been found (Esfeld et al. 2018) and it has been observed that a reversal to a lost ancestral state could happen through changes in relatively few genes (Seher et al. 2012). Moreover, some observations suggest that the molecular blueprint associated to a trait could be preserved even in its phenotypical absence (*e*.*g*. Marshal et al. 1994; Carlini et al. 2013; Leal and Cohn, 2016), possibily due to pleiotropic constrains (Lammers et al. 2019).

Among the many challenges to Dollo’s law, the evolution of wings in phasmids stands as one of the first and more iconic examples (Whiting et al. 2003). It also played a central role in rethinking Dollo’s law and in initiating a long-standing debate on the topic (Collin and Miglietta 2008). This polyneopteran order of insects includes over 450 genera and 3300 described species, out of which 40% *circa* are macropterous while the remaining are either brachypterous or apterous. As can be observed in Figure 1, phasmids’ wings present a high level of anatomical disparity and differences can be found at all taxonomic levels, including between congeneric species (Zeng et al. 2020). Macropterous phasmid species present long wings which are able to sustain flight with different efficiency, depending on the wing to body size ratio; however, most of them are considered weak flyers, using wings mainly for controlling free-fall descents from tree canopies (Maginnis 2006). Some lineages have also evolved non-aerodynamic functions for wings, such as aposematic coloration or stridulation capacity, typical of brachypterous forms.

**Figure 1.**
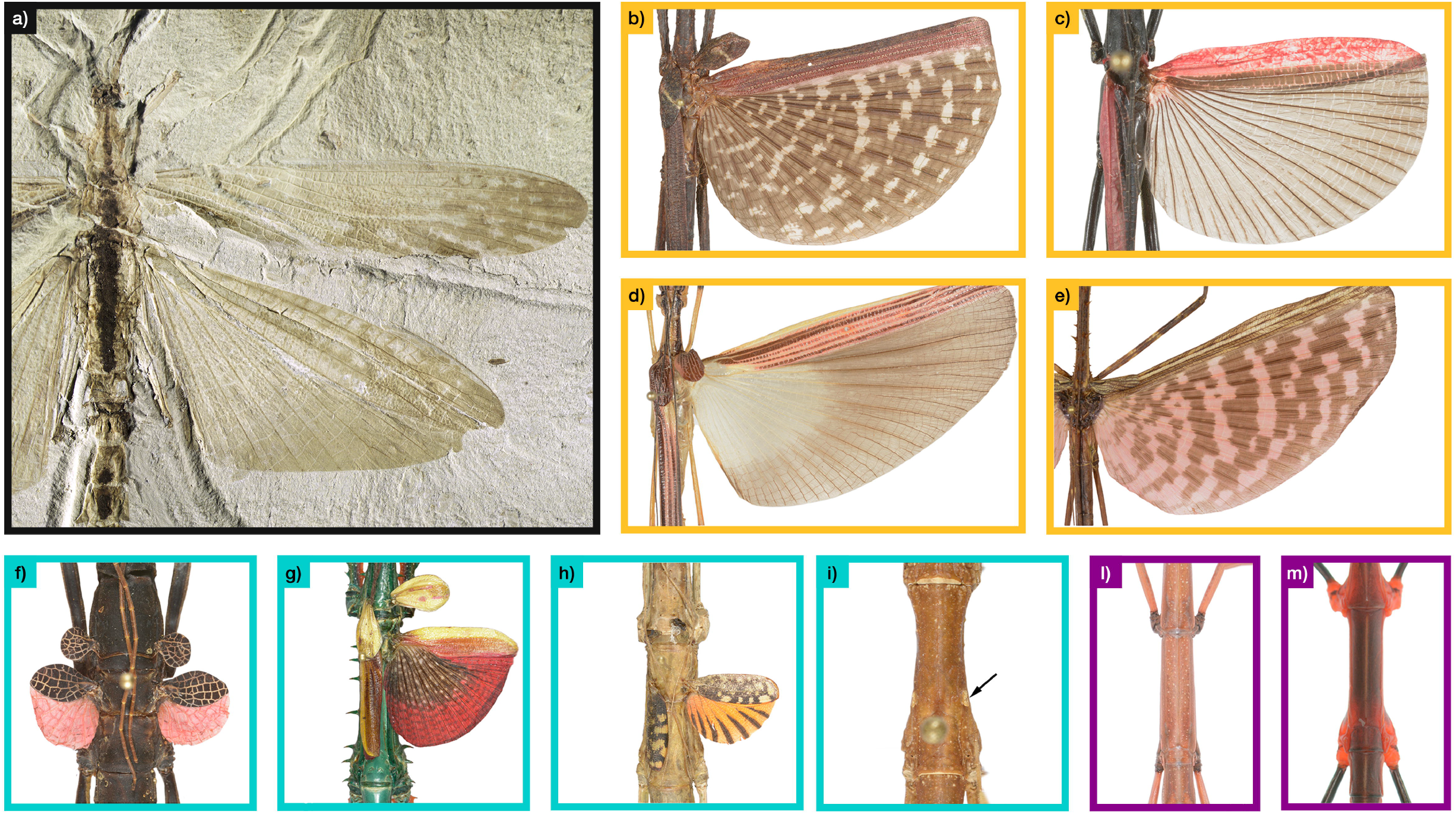
Wings disparity among extinct and extant Phasmatodea. (a) *Aclistophasma echinulatum* Yang, Shi, Engel, Zhao, Ren and Gao, 2021; (b) *Pterinoxylus crassus* Kirby, 1889; (c) *Orthomeria kangi* Vallotto, Bresseel, Heitzmann and Gottardo, 2016; (d) *Parastratocles fuscomarginatus* Conle, Hennemann, Bellanger, Lelong, Jourdan and Valero, 2020; (e) *Diesbachia tamyris* (Westwood, 1859); (f) *Peruphasma schultei* Conle and Hennemann, 2005; (g) *Achrioptera manga* Glaw, Hawlitschek, Dunz, Goldberg and Bradler, 2019; (h) *Phaenopharos struthioneus* (Westwood, 1859); (i) *Hypocyrtus ornatissimus* (Brunner von Wattenwyl, 1907); (l) *Bacillus atticus* Brunner von Wattenwyl, 1882; (m) *Oreophoetes peruana* (Saussure, 1868). Boxes are colored correspondingly to trait state: gold = Macroptery, turquoise = Brachyptery, violet = Aptery. All photographs have been taken by Pablo Valero, with the exception of *A. echinulatum* which has been provided by Hongru Yang.

Wings emerged early in the diversification of insects with frequent losses occurring during their evolutionary history (Wipfler et al. 2019). In 2003, Whiting et al. proposed that wings disappeared in the lineage that led to extant phasmids and subsequent reversals restored the lost structures throughout the clade evolutionary history. Shortly after this claim several comments followed (Stone and French 2003; Trueman et al. 2004) and nowadays - even if there is a growing consensus that lost traits can be re-acquired (Porter and Crandall 2003; Wiens 2011) - several possible weak spots have been exposed in the paper by Whiting et al. and many approaches have been proposed to avoid incorrect rejection of Dollo’s law. Root prior probability (Sauquet et al. 2017; Goldberg and Igi• 2008), trait-dependent diversification rates (Goldberg and Igi• 2008; Holland et al. 2020) and tree uncertainty (Bollback 2006; Rangel et al. 2015) have all to be carefully considered when testing for the irreversible evolution of a trait.

Phylogenetic tests of trait irreversible evolution are frequently misled by inappropriate assignment of the character state distribution at the root, which should be avoided unless unequivocal evidences are available. Despite phylogenetic analyses support extinct Phasmatodea clades - such as Gallophasmatidae, Pterophasmatidae and Susumanioidea - as stem groups of modern stick and leaf insects, they don’t allow to make conclusive considerations of the wing status of the MRCA of the extant species (Yang et al. 2019; Yang et al. 2021). Nonetheless, a stark difference can be observed between the wings of extinct stem phasmids and extant species: the latter have either reduced or absent tegmina while stem fossils specimens present both pairs of wings full-sized. This is the case of the fossil specimen *Aclistophasma echinulatum*, which is found in Figure 1a. In insects, wings confer a plethora of potential advantages, such as evading predators, dispersal and mate-finding (Goldsworthy 2018), but partial reduction or complete loss can have adaptive value as well: wings loss has been related to increased female fecundity (Roff 1994), cryptic capacity (Whiting et al. 2003) and also to trade-offs in resource allocation between different anatomical structures (Maginnis 2006). Wings can be associated with sexual recognition and selection, which can drive diversification dynamics (Arnqvist et al. 2000; Masta and Maddison 2002; Singh and Singh 2014) and different flight capabilities are associated to differential diversification rates in several insect clades, with both dispersal reduction and increment acting as possible drivers of diversification (Ikeda et al. 2012; Misof et al. 2014; Waters et al. 2020;). When the state of a character affects the diversification rates of a lineage, it can ultimately also drive the trait distribution along the phylogeny tips; this phenomenon can lead to strong biases in transition rates and ancestral state reconstruction when common MK models - which assume neutral character evolution - are used (Goldberg and Igi• 2008; Holland et al. 2020). Trait subject to Dollo’s law are expected to become less frequent throughout the evolution of a lineage, but they can be instead rather widespread if they can drive diversification dynamics of clades. Thus, the interplay between wings and phasmids diversification rates needs to be carefully considered when testing for the trait irreversible evolution. An additional obstacle in elucidating wings evolutionary patterns comes from the phylogenetic uncertainty which is often associated with Euphasmatodea (*i*.*e*. all Phasmatodea excluding *Timema*). While a recent transcriptomic sequencing effort represents fundamental progresses for understanding the systematic relationships of Euphasmatodea major clades (Simon et al. 2019; further refined in Tihelka et al. 2020), the clade still present uncertain instances in both systematic relationships and divergence times (*e*.*g*. Robertson et al. 2018). As such, this kind of uncertainty cannot be disregarded when studying macroevolutionary patterns in the clade.

In this study we reconstructed the most comprehensive time-tree so far for phasmids and leveraged it to explore the evolutionary patterns of wings in the clade. We tested the hypothesis that reversals could restore a trait to its former state subsequently to its loss by taking advantage of multiple approaches, including different trait-coding strategies along with global (model-fitting) and local (ancestral state reconstruction and stochastic character mapping) approaches. We took into account common biases in phylogenetic comparative analyses and implemented all possible recommendations proposed for testing Dollo’s law rejection, such as trait-dependent diversification rates and phylogenetic uncertainty. Furthermore, we tested the impact on our findings of different assumptions about the probability of wings losses and reversals. By addressing these questions relatively to phasmid’s wings, we used an iconic case to test to which extent the evolution of a complex trait can be a reversible process and to widen the knowledge on how complex phenotypes evolve.

## Material and Methods

### Phylogenetics dataset

Sample were collected, morphologically identified and preserved dry or in ethanol until molecular analyses were carried out. Genomic DNA was extracted from leg tissues of 111 individuals (Supplementary Table 1), using the protocol of “Smarter Nucleic Acid Preparation” (Stratec). Eight molecular markers were amplified, consisting in: two mitochondrial (two fragments of the Cytochrome Oxidase subunit I; one of the Cytochrome Oxidase subunit II) and one nuclear (Histone subunit 3) protein coding genes (PCGs) along with two mitochondrial (12S; 16S) and two nuclear (28S; 18S) ribosomal RNAs (rRNAs). PCR reactions were carried out according to standard protocols; primers sequences and annealing conditions can be found in Supplementary Table 2. PCR products were screened through 1% agarose gel electrophoresis and purified using ExoSAP PCR Product Cleanup Reagent (Thermofisher). Amplicons were Sanger sequenced by Macrogen Europe Lab. Chromatograms were inspected with SeqTrace 0.9.0 (Stucky 2012) and the resulting sequences were visualized using Aliview v1.26 (Larsson 2014). BLASTn (Zhang 1997) on NCBI Genbank database was used to search for potential contaminants. Complementary sequences were obtained from unpublished inhouse projects on Euphasmatodea systematics (Genbank accession numbers MN449491-MN449962 and MT077516-MT077845) and from all specimens for which a species level identification was available from the following papers: Whiting et al. (2003), Buckley et al. (2009), Buckley et al. (2010), Bradler et al. (2014), Bradler et al. (2015), Goldberg et al. (2015), Robertson et al. (2018) and Glaw et al. (2019) plus outgroup sequences belonging to Notoptera (Grylloblattodea + Mantophasmatodea; Damgaard et al. 2008; Jarvis and Whiting 2006) (Supplementary Table 1). Taxonomical annotation is provided in Supplementary Table 1. We did not consider Embioptera - which would represent the true sister group of Phasmatodea - as both previously published (Song et al. 2016) and our own exploratory analyses showed that their inclusion introduces long-branch attraction artifacts.

### Rogue taxa removal and time-tree inference

All sequences were aligned using Mafft 7.402 (Katoh and Standley 2013): PCGs were translated to amino-acids using AliView v1.26, aligned using the “L-INS-i” algorithm and subsequently retro-translated into nucleotides. rRNAs were aligned using the “--X-INS-i” algorithm to take into account their secondary structure. Gblock v. 0.91b (Castresana 2000) was used to exclude possible misaligned positions, selecting the codon flag for PCGs and the nucleotide one for rRNAs. Others parameters were set as follow: minimum number of sequences for a conserved position, 50% (PCGs and rRNA) of sequences included in the alignment; minimum number of sequences for a flanking position, 70% for PCGs and 60% for rRNAs; maximum number of contiguous no conserved position, 8 for PCGs and 10 for rRNAs; minimum length of a block, 10 for PCGs and 5 for rRNAs; allowed gap position, *all* for PCGs and *with half* for rRNAs. Each MSAs has been then concatenated using Phyutility v. 2.2 (Smith and Dunn 2008). All subsequent phylogenetic inferences were performed on the Cipres Science Gateway (http://www.phylo.org/portal2), using XSEDE (Miller et al. 2010).

Maximum Likelihood (ML) inference was performed using IQ-TREE 1.6.12 (Nguyen et al. 2015), the concatenated alignment was partitioned *a priori* by gene and codon position. The best-fit partitioning scheme and evolutionary model were chosen using ModelFinder (Kalyaanamoorthy et al. 2017) according to the BIC score (Supplementary Table 3). Branch support was estimated with 1000 UltraFast parametric bootstrap replicates (Minh et al. 2013). RogueNaRok (Aberer et al. 2013) was used to identify and remove rough taxa until none was found.

We carried out two Bayesian Inferences (BI) with Beast2 (Bouckaert et al. 2014): a first one using only the monophyletic constrains associated to fossil calibrations (Supplementary Table 4) and a second one including additional monophyletic constrains to match the phylogenomic resolution of Tihelka et al. (Supplementary Materials). In both instances, divergence times estimation, tree inference and model averaging were jointly performed with the bModelTest package (Bouckaert and Drummond 2017), using a fully Bayesian framework. The concatenated alignment was partitioned *a priori* by gene with unlinked site models and linked tree and clock models; a relaxed clock with a lognormal distribution and a Birth-Death model were used as clock and tree priors, respectively. An exponential distribution was chosen as prior distribution with a minimum hard bound set up to the age of the fossil and a soft maximum bound of 410 Ma. Two independent chains were run for 400 million generations and sampled every 5000 states. After convergence and adequate ESS were assessed (> 200) with Tracer v.1.7.1 (Rambaut et al. 2018), log files and tree files were combined with LogCombiner v. 2.6.2 and a 25% conservative burn-in was For subsequent macroevolutionary analyses, we used: (a) a random sampling of 1000 trees from the posterior distribution of the BI inference which leveraged only the monophyletic constrains associated to fossil calibrations (from here on UnConstrained Trees or UC-Trees) or (b) the maximum clade credibility tree derived from the BI which leverages phylogenomic constrains (from here on Phylogenomic Constrained Tree or PC-Tree); for the latter, trees were summarized keeping the median node heights in TreeAnnotator v. 2.4.2. Analyses using UC-Trees allowed us to take into consideration the impact of uncertainty in systematic relationships and divergence times while analyses on the PC-Tree leveraged the current best single hypothesis for the evolutionary history of Euphasmatodea.

### Morphological datasets

The morphological datasets of wings states were compiled from all available sources, including collected specimens and images on the Phasmida Species File portal (http://phasmida.speciesfile.org; last accessed in July 2020). We considered wings as a species-level trait and in the case of sexual dimorphism the more “complex” structure between the two sexes was considered (*e*.*g*. a species with apterous females and brachypterous males was considered as brachypterous). In a first coding scheme, wing morphology was coded as a 2-states trait, with presence as 1 (including both macroptery and brachyptery) and absence as 0. An alternative 3-states coding scheme considered macroptery and brachyptery as different states, so that 0: apterous, 1: brachypterous and 2: macropterous species. Brachypterous species were defined as having alae reaching up to the posterior margin of abdominal tergum II, while we considered as macropterous species having alae projecting over the posterior margin of tergum II. All coding schemes are provided in Supplementary Table 1.

For both the PC-Tree and the UC-Trees, we pruned all outgroup species and a single ingroup species, *Dimorphodes prostasis* Westwood, 1859, for which no conclusive morphological information was available.

### Comparative analyses considering two states (winged, apterous)

We initially used a model comparison approach, evaluating three common MK models through the *fitDiscrete* function of the R package geiger v2.0 (Pennell et al. 2014). These were: (1) Equal Rates (ER), (2) All Rates Different (ARD) and (3) Loss Only (LO). All the models were fitted on the PC-Tree and compared looking at the resulting AICc score. To take into account tree uncertainty in model selection and parameters estimations we used the *influ_discrete* function from the R package sensiPhy v0.8.5 (Paterno et al. 2018), fitting all aforementioned models to the 1000 UC-Trees.

Ancestral state reconstructions (ASRs) were performed on the PC-Tree with the best-fit model of evolution and the other models. The *rayDISC* function and a prior root probability weighted accordingly to the method of Maddison et al. (2007) and FitzJohn et al. (2009) were used, so that root states probability is weighted according to their conditional probability given the data. As this assumption can greatly influence ASR and parameters estimation (Goldberg and Igi• 2008; Sauquet et al. 2017) we performed two additional ASRs with ER and ARD models and a flat prior, so that root state is equally likely to be in state 0 and 1.

To take into account parameter and tree uncertainty in ASR we performed stochastic character mapping (SCM) using the *make*.*simmap* function, implemented in Phytools (Revell 2012), under the best-fit model on the PC-Tree with 100 simulation (100Sim) and across 100 randomly-sampled UC-Trees with one simulation (100Trees).

### Comparative analyses considering three states (macropterous, brachypterous, apterous)

When considering brachyptery, our comparative analyses followed the same approach used with the 2-states character coding scheme, but taking into account a wider range of evolutionary models. We fitted the three unordered MK models previously used: (1) ER, (2) ARD, (3) LO, adding a (4) symmetrical model (SYM). We also took into consideration some additional biologically meaningful user-supplied MK models, in which one or more transitions were fixed to 0: (5) Ordered Loss Only (O-LO) in which only losses between proximal states were allowed; (6) Partial Reversal Brachyptery (PR-BP) where all losses and transitions from brachypterous to macropterous forms were allowed; (7) Partial Reversal Aptery Brachiptery (PR-AB) where all losses and transitions from apterous to brachypterous were allowed; (8) Ordered Complete Gain (O-CG), with all losses allowed plus transitions from apterous to brachypterous and from brachypterous to macropterous; (9) Ordered Full Model (O-FM), where only transitions between proximal states were allowed. As before, this model fitting approach was carried out both on the PC-Tree and the 1000 randomly-sampled UC-Trees. In the latter case, for simplicity and due to computational limits, we excluded from these analyses the O-LO and PR-AB models (see Results for justification). As sensiPhy can handle only binary characters, we used a custom script to run the *fitDiscrete* function on the 1000 UC-Trees.

As the outcome of model-fitting analyses was quite conclusive (see Results), we conducted the ASRs on the PC-Tree using *rayDISC* with the best-fit model only and a “maddfitz” or a “flat” prior. Since we noticed that - differently from the analyses carried on with the 2-states dataset - the transition rates inferred by *rayDISC* and *fitDiscrete* were different, we also carried out an additional ASR to mirror previous model selection analyses with *fitDiscrete* (forcing transition rates calculated by *fitDiscrete* and a “maddfitz” root prior).

As for the binary dataset, tree and parameter uncertainty in ASR was taken into consideration carrying out two SCM analyses respectively with 100 simulations on the PC-Tree (100Sim) and 100 randomly sampled UC-Trees with one simulation (100Trees).

### Trait-dependent diversification analyses

To assess if trait-dependent diversification rates could bias our previous analyses, we relied on the Hidden State Speciation and Extinction framework (HiSSE) in R (Beaulieu and O’Meara 2016). The association between shifts in phasmids diversification rates and the evolution of wings and flight capabilities has been tested using two distinct character coding schemes, respectively: (a) the one used in previous 2-states analyses (macroptery and brachyptery as 1; aptery as 0) and (b) a modified version with macroptery coded as 1 and brachyptery and aptery as 0 (also including four species which are macropterous but not able to fly due to their excessive size; Supplementary Table 1 and Supplementary Materials). We tested a wide range of possible macro-evolutionary scenarios on the PC-Tree, generating several state-dependent diversification models in a HiSSE /BiSSE framework: (1) BiSSE: a standard BiSSE model where diversification rates depend on the wings state; (2) Full HiSSE: diversification rates depend on one explicit character (wings) and a hidden one, differently for winged and apterous forms (*i*.*e*. a common full HiSSE model with the hidden states A and B); (3) HiSSE: diversification rates change between apterous and winged species only for one hidden state (in our case B, but is arbitrary since they are unknown), while for the other (A in our case) they result equal; (4) and (5) Half HiSSE 1 and Half HiSSE 2: diversification rates depend on the explicit character and on a hidden state but only for respectively winged and apterous species. These models were tested against three null models: (6) BiSSE Null: diversification rates constant through the tree and no hidden states; (7) CID-2: diversification rates change only as a function of two hidden state; (8) CID-4: a null model in which diversification rates are independent from the explicit trait but depend only on four hidden states, accounting for the same number of diversification parameters of a full HiSSE model (*i*.*e*. eight). The CID-2 and CID-4 model has been proposed to be two good null hypotheses in order to avoid common Type I errors (Beaulieu and O’Meara 2016). As for previous analyses, all the models were compared through AICc scores.

### Impact on wings reversals inference of different assumption on their probability

As previous approaches leveraged no information on transitions rates between different states other than the trait distribution on the tips of the tree and the tree itself, we explored the impact of different assumptions of the probability of wings reversals on our analyses. We assumed a diminishing probability of reversals compared to loss (from 1:1 to 1:500 with an interval of 10, corresponding to an ER model and an approximation of a LO one) in conjunction with loss rates between 0.001 and 0.02, with an interval of 0.001. These rates have been chosen to reflect the optimized values found in previous analyses (0.00350 for the ER model and 0.00886for the LO; see Results). We carried out SCM using Phytools function using either 100 simulation on the PC-Tree (100Sim) and 100 random UC-Trees with one simulation (100Trees) for each combination of reversals relative probability and model rate.

## Results

### Datasets and time-tree inference

The concatenated alignment resulted in 4112 positions and included 345 taxa, 322 of which were phasmids (Supplementary Materials). Out of the 322 phasmid taxa, 92 represent new specimens and 230 were obtained from NCBI. We were able to obtain information about wing state for all Phasmatodea species with the exception of *Dimorphodes prostasis* Westwood, 1859, which has been excluded from comparative analyses. Overall, 180 species were identified as apterous (56%), 33 as brachypterous (10%) and 108 as macropterous (34%) (Supplementary Table 1). The 1000 randomly sampled UC-Trees presented a wide range of arrangements among phasmids major clades, yet major and established phasmids clades are retrieved as monophyletic (Supplementary Materials). The constrained PC-Tree (Supplementary Figure 1) recovered the divergence between the two suborders Timematodea and Euphasmatodea at 180 million years ago (Ma) (95% HPD = 122-249 Ma), with the Euphasmatodea radiation beginning 108 Ma (95% HPD = 75-148 Ma) and the diversification of major lineages happening in the subsequent 30 million years.

### Comparative analyses considering two states (winged, apterous)

For the 2-states character coding scheme, the best-fit on the PC-Tree resulted to be the ER model (AICc = 276.5326, Supplementary Table 5A) with a transition rate between the two states equal to 0.00350 (Supplementary Figure 2A), followed by the ARD (AICc = 278.1931), with transition rates similar to those of the ER model. LO model had a •AICc = 28.894.

Under the best-fit model, the root of the PC-Tree was reconstructed as apterous with the maximum posterior probability (PP). Nine ancient (*i*.*e*. along internal branches) and fourteen recent (*i*.*e*. along terminal branches) reversals of wings were inferred, while wing losses were inferred respectively five and ten times (total number of transitions: 38; Figure 2A). Similar results were obtained when the ASR was carried out under the ARD model, due to the similar inferred transition rates (Supplementary Figure 3). Instead, in a scenario consistent with Dollo’s law, 61 independent losses (34 recent and 27 ancient) of wings were necessary to explain the distribution of the trait throughout the tree (Supplementary Figure 3). Even using a flat prior, the root was recovered as apterous for both ER and ARD models (PP > 75) with transition rates, AICc values and the overall ASRs approximately equal to those obtained in previous analyses (Supplementary Table 6A, Supplementary Figure 3).

**Figure 2.**
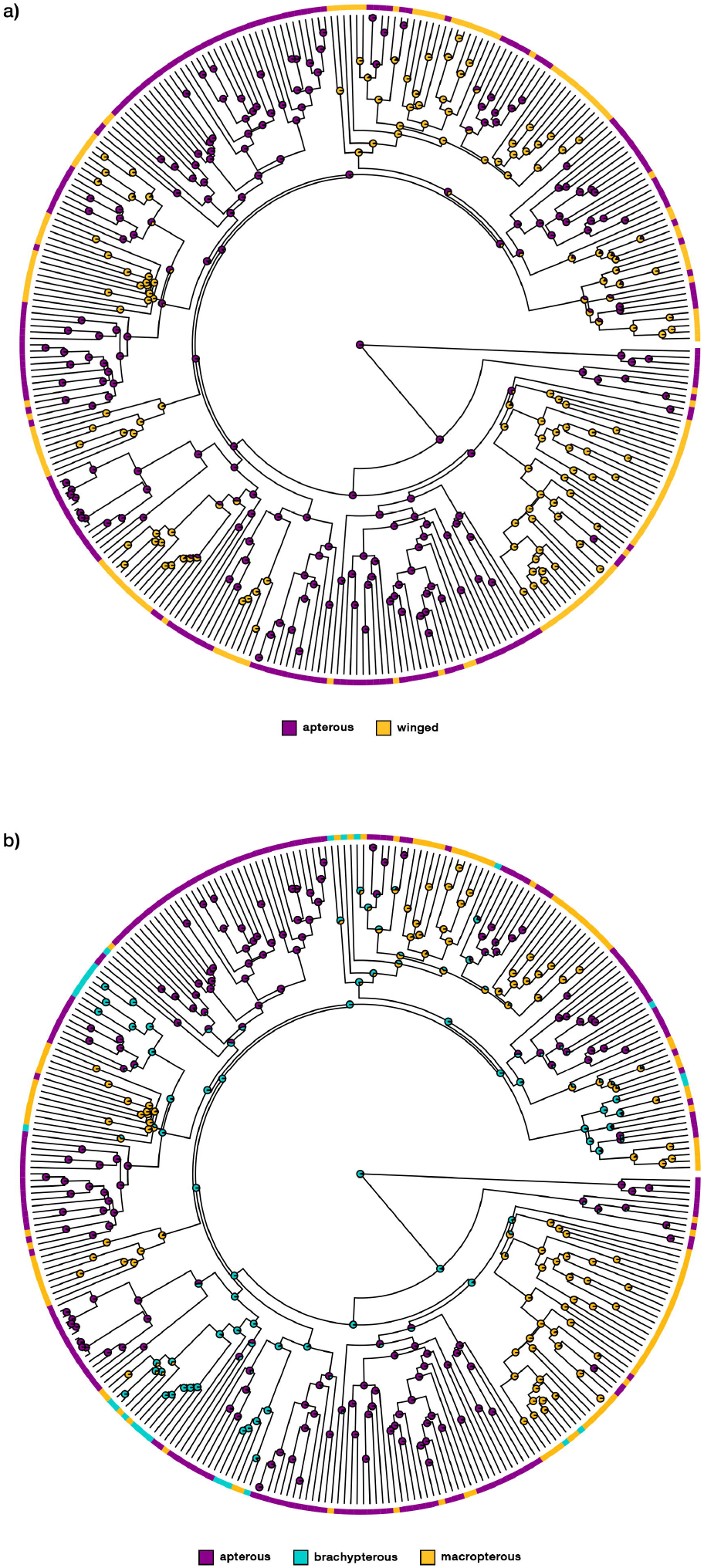
Phasmatodea wings ancestral state reconstruction on the maximum clade credibility tree obtained from the Bayesian Inference which includes monophyletic constrains to match the phylogenomic resolution of Tihelka et al. (2020) (PC-Tree); (a) and (b) leverage 2-states and 3-states coding schemes under the best-fit model of trait evolution, respectively Equal Rates with a maddfitz root prior and All Rates Different with a maddfitz root prior and fixed transition rates.

Sensitivity analyses highlighted a strong preference for the ER model (Figure 3A): it was recovered as the best-fit for 993 trees out of the 1000 taken from the BEAST posterior distribution, and its AICc distribution resulted significantly lower than all the others (Kruskal-Wallis test, p < 0.001; post-hoc pairwise Wilcoxon test with Bonferroni correction, p < 0.001). For the remaining 7 trees, the ARD model was found the best-fitting.

**Figure 3.**
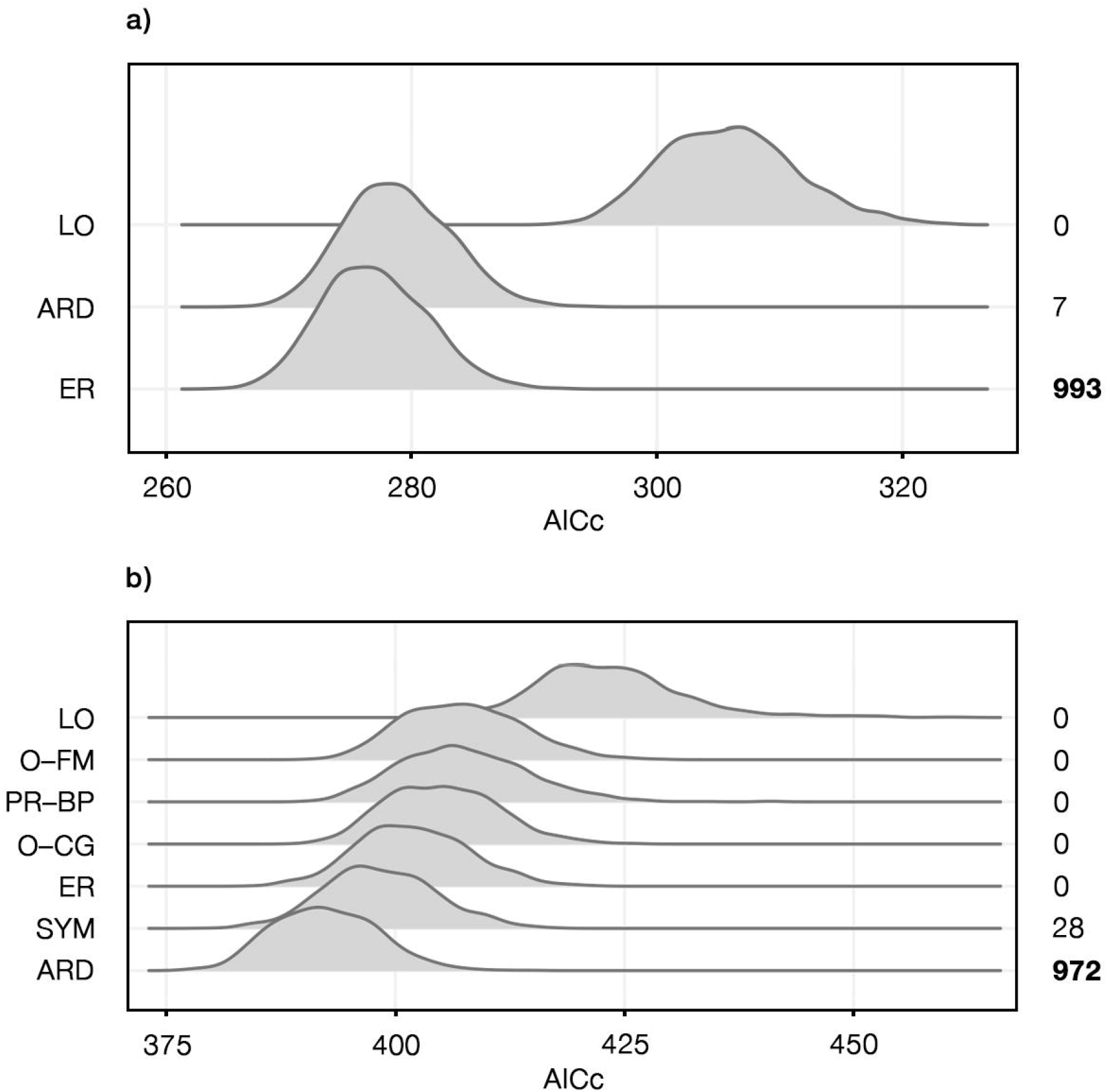
Model-fitting considering uncertainty in tree topology and branch lengths. Analyses have been performed on the 2-states (a) and 3-states (b) character coding schemes for 1000 trees randomly sampled from the posterior distribution of the “unconstrained” Bayesian Inference (UC-Trees). On the right side the number of trees for which each model resulted to be the best-fit is reported, with the more frequent ones (respectively ER and ARD) highlighted in bold. Models are: LO = Loss Only; O-FM = Ordered Full Model, where only transitions between proximal states were allowed; PR-BP = Partial Reversal Brachyptery, where all losses and transitions from brachypterous to macropterous forms were allowed; O-CG = Ordered Complete Gain, with all losses allowed plus transitions from apterous to brachypterous and from brachypterous to macropterous; ER = Equal Rates; ARD = All Rates Different.

Stochastic character mapping highlights a weak impact of parameter, topology and branch lengths: both 100Sim and 100Trees analyses recovered a high preference for a apterous root (Figure 4A) and a higher number of reversals than losses (Figure 4B). An average of 46.27 and 48.01 changes between states were inferred respectively for the 100Sim and 100Tree, with an average of 27.23 and 27.14 being reversals to winged forms.

**Figure 4.**
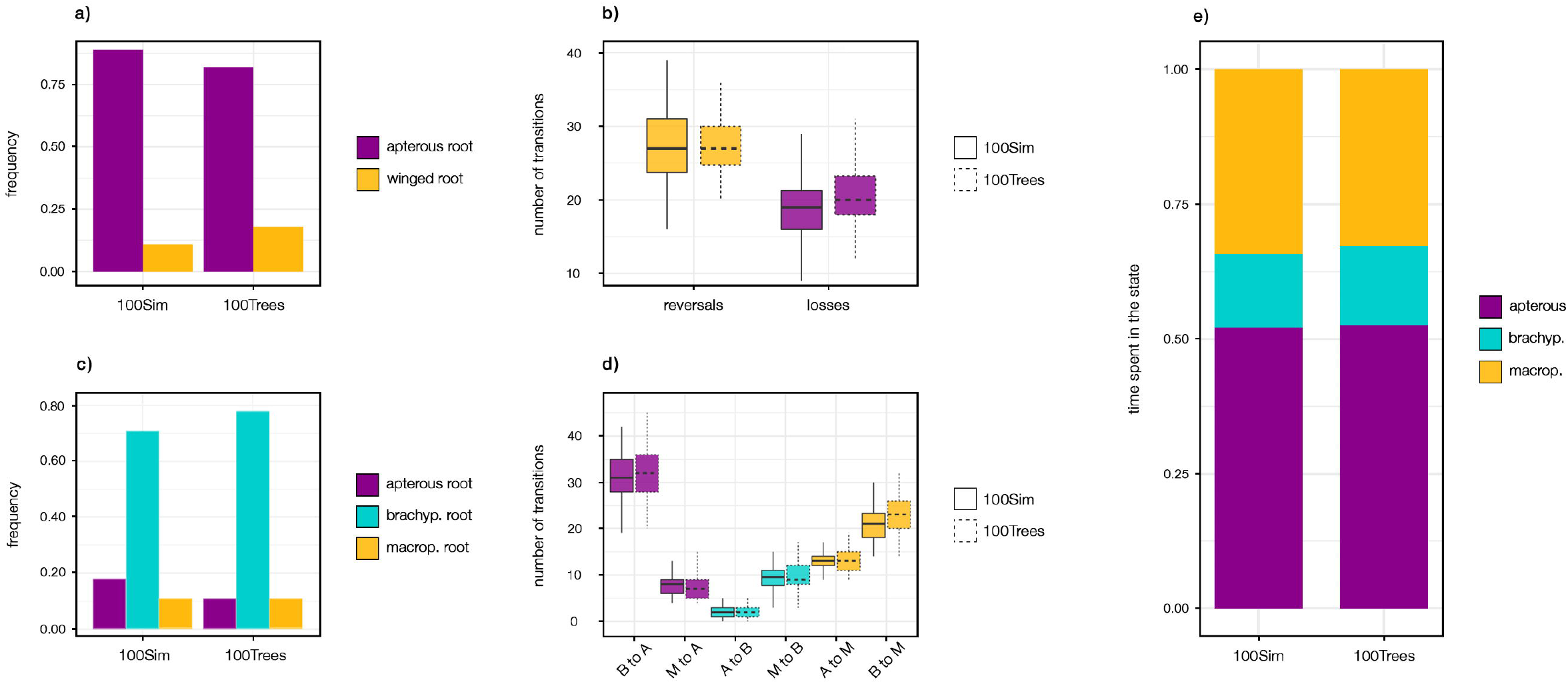
Stochastic character mapping leveraging 2-states and 3-states coding schemes: the frequency of root state is represented in (a) and (c). (b) and (d) describe the average number of inferred transitions. (e) represents the time spent in each state for the 3-states character coding scheme. In (b) and (d) boxplots are colored correspondingly to transitions end state, so that: gold = Macroptery, turquoise = Brachyptery, violet = Aptery. 100Sim refers to Stochastic Character Mapping analyses with 100 simulations on the PC-Tree; 100Trees refers to the same analyses carried on 100 randomly-sampled UC-Trees with a single simulation.

### Comparative analyses considering three states (macropterous, brachypterous, apterous)

When implementing brachyptery as a third state, ARD resulted the best-fitting model (AICc= 389.5214) on the Bayesian PC-Tree, followed by SYM (•AICc = 3.744) and ER (•AICc = 6.597) (Supplementary Table 5B). Again, LO model was rejected with a •AICc = 33.12. The two models O-LO and PR-AB resulted as the less-supported, with a •AICc of 33.671 and 43.408, respectively; therefore, we discarded them from subsequent analyses. Transition rates of the ARD model are shown in Supplementary Figure 2B. The highest transition rates are those which describe the shifts from brachyptery to either macropterous (0.01117) or apterous (0.0169) forms, while the rates from apterous to fully macropterous forms and *vice-versa* resulted to be closely matching (0.00181 and 0.00166).

The ASRs carried out on the Bayesian PC-Tree under the ARD model resulted in different outcomes depending on the parameters used (Supplementary Figure 5), but in all reconstruction reversal to macropterous or brachypterous forms from apterous ancestors are inferred. Using a flat root prior and a maddfitz one with fixed rates, the root was reconstructed as brachypterous with a PP of 0.66 and 0.91, respectively (Supplementary Table 6B). In these two ASRs wings reversal from apterous to brachypterous or macropterous forms are present, but only as recent events on terminal branches. Though, a large number of transitions from brachyptery to both macroptery and aptery are recovered, reflecting the low rates between apterous and macropterous forms and the high rates that move away from partially developed wings. Using a maddfitz prior with transition rates inferred directly by *rayDISC*, the root is recovered as apterous with a PP of 0.74 and reversals are inferred also throughout internal branches. These results reflect the different transition rates inferred with respect to previous analyses: the lowest transition rate was found between brachypterous and winged forms, while higher rates were inferred towards brachyptery from both apterous and macropterous forms. (Supplementary Table 6B). As such, this model favoured an apterous state for deepest nodes and more transitions to brachyptery (Supplementary Figure 5). However, the two ASRs which inferred the root as brachypterous resulted as better supported than those inferring an apterous one (•AICc = 6.94), with a slight preference for the one obtained with fixed parameters (•AICc = 0.85; Supplementary Table 6B).

Sensitivity analyses highlight a strong preference of the ARD model which was recovered as the best-fit for 972 of the 1000 randomly sampled UC-Trees, with a AICc distribution significantly lower with respect to all others (Kruskal-Wallis test, p < 0.001; post-hoc pairwise Wilcoxon test with Bonferroni correction, p < 0.001), while the SYM model was preferred for the remaining 28 trees (Figure 3B).

Regarding SCMs, a brachypterous root was preferred in both analyses (Figure 4C) and an average of 84.79 and 87.76 changes between states were inferred for the 100Sim and the 100Trees analyses, respectively. In both of them, the highest average number of transitions were recovered to be from brachypterous to apterous forms (31.19 for the 100Sim and 31.94 for the 100Trees), followed by transition from brachypterous to fully macropterous (respectively 21.05 and 23.05) (Figure 4D). The same results are reflected by the mean total time spent in each state, with brachyptery being the less represented (14% for both analyses; Figure 4E).

### Trait-dependent diversification analyses

All our diversification analyses favoured a null CID-2 model, implying a decoupling of phasmids diversification dynamics with their wings development and flight ability. Indeed, both full Hisse and Bisse models, which imply variations in diversification rates according to trait state, resulted to have a lower support, with a minimum •AICc of respectively 8 and 75. (Supplementary Tables 7A – 7B).

### Impact on wings reversals inference of different assumption on their probability

When different assumptions on the relative probability of reversals (ratio of wings reversal to loss rates) are tested in a SCM framework, it can be observed that up to a ratio of 1:30 reversals are consistently inferred in both the 100Sim and 100Trees analyses (at least one reversal is inferred in each of the 100 SCM analyses) independently from the absolute values of the evolutionary model parameters (Figure 5; Supplementary Figure 5). The latter strongly affect the consistency of reversals inference when the ratio of reversal to loss is assumed lower than 1:30: at low model rates, reversals are inferred even when assumed to be five hundred times less likely than losses. Remarkably, no evolutionary model considered in this analysis consistently rules out the possibility of a reversal, which are always inferred under certain tree and model parameter conditions.

**Figure 5.**
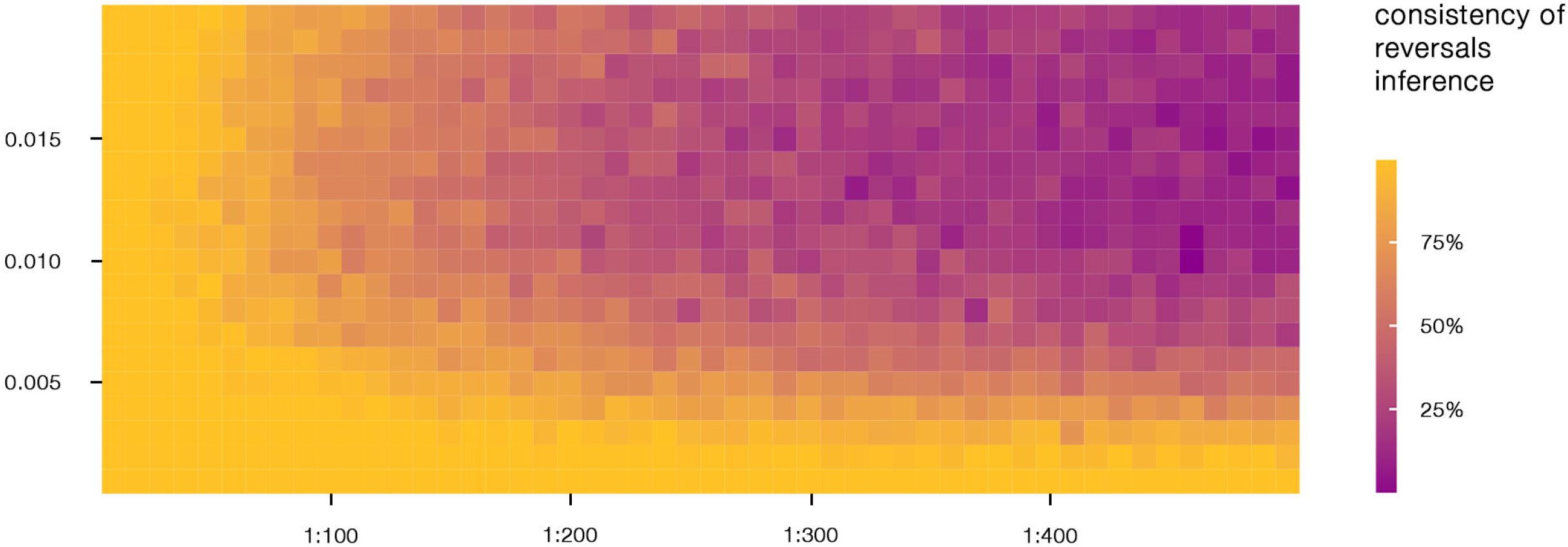
Impact on wings reversal inference of different assumption on their probability. Tile coloring represents the consistency of reversals inference on the PC-Tree when different model rates (x-axis) and relative probability of reversal (y-axis) combinations are assumed. When consistency equals 100%, at least one reversal is inferred in each of the 100 simulations of the stochastic character mapping analysis performed using the relative model parameters combination; if consistency <100% and >0% reversals are inferred just on a fraction of the considered trees and character histories.

## Discussion

Our phylogenetic reconstruction of Phasmatodea evolution represents the most comprehensive one to date. Yet, our phylogenetic inferences highlighted once more the difficulties in finding a stable and supported backbone for Euphasmatodea phylogeny using a handful of gene fragments, due to the scarce phylogenetic signal associated to their rapid evolutionary radiation (*e*.*g*. Robertson et al. 2018; Glaw et al. 2019). Moreover, in phasmids, uncertainty is not just limited to systematic relationships but it is also found in the timing of their evolution. In previous studies, the split between Euphasmatodea and *Timema* ranged between ∼95 Ma and ∼270 Ma (Buckley et al. 2009; Forni et al. 2021), while our estimate set it at ∼174.5 Ma (95% HPD: 122.63-249.1 Ma); this date falls between those inferred in the recent phylogenomic efforts by Simon et al. (2019) and Tihelka et al. (2020) (respectively ∼121 and ∼250 Ma). The same holds for the Euphasmatodea crown node, which is here retrieved in the Mid-Cretaceous at ∼105 Ma (95% HPD: 75-148 Ma), while in phylogenomic analyses ranges from ∼80 to ∼150 Ma. Such striking incongruences are also found at a lower taxonomic level (Bank et al 2021b) and can be largely explained by the different choice and /or usage of fossil calibration; our approach has been to include in our analyses only fossils which are deemed uncontroversial in their placement (Bradler and Buckley, 2011) and to implement wide upper boundaries of prior distribution in order not to force any hypothesis. Paleontological evidence (Yang et al. 2019) are consistent with our results, indicating the Mid-Cretaceous as an important period of phasmids diversification, possibly related to the contemporary rapid diversification of angiosperms (Peris et al. 2017). Throughout this paper we carefully considered phasmids uncertainty in systematic relationship and divergence times by leveraging two kind of analyses: one relying on the single best phylogenetic hypothesis currently available (PC-Tree) and another which allowed us to consider extensively uncertainty in topology and branch lengths (UC-Trees).

A first hint of the reversible evolution of phasmids wings is simply provided by the distribution of the trait states across the phylogeny (Figure 2). Although many apterous, brachypterous and macropterous species form well supported clades, other include a mixture of different states: for example, several macropterous species, like *Bacteria ploiaria* (Westwood, 1859), *Cranidium gibbosum* (Burmeister, 1838) and *Lobofemora bidoupensis* (Bresseel and Constant, 2015), are found in clades mainly including apterous taxa. This is also the case of some winged Heteropterygidae species which - as recently corroborated by Bank et al. (2021a) - are found nested in a largely brachypterous / apterous clade. Nonetheless, this kind of pattern appears to be consistent with shallow morphological taxonomy and it has to be considered that we carefully excluded any rogue taxa from our analyses.

Common MK models assume neutral character evolution and systematic biases can arise when a trait has the potential to influence probabilities of lineage speciation and extinction. *SSE methods can outperform common MK models in state-dependent diversifications scenarios, allowing parameters to depend on the state of the character (Goldberg and Igi• 2008; Holland et al. 2020). Trait-dependent diversification analyses presented here reject the hypothesis that wing development and flight ability are associated with shifts in diversification rates in phasmids. This result contrasts with the earlier hypothesis that wings loss is correlated to increased speciation rates in stick insects (Zeng et al. 2020) and allowed us to exclude this potential bias while testing for Dollo’s law rejection.

Model selection strongly supported models in which reversals occurred (*i*.*e*. from apterous to winged forms in the 2-states analyses; from apterous to bachypterous and macropterous forms and /or from brachypterous to macropterous ones in the 3-states analyses). These outcomes were consistent for both the PC-Tree and the UC-Trees analyses, while a model consistent with Dollo’s law - *e*.*g*. LO - was never supported. Interestingly, brachyptery resulted to be the most unstable state, showing the highest rates of transition departing from it compared to other states. Similar outcomes are provided by SCM analyses (both 100Sim and 100Trees): reversals are consistently inferred and brachyptery is recovered as the state with the greatest number of transitions moving away from it and with the least time spent within, (Figure 4). Altogether, these evidences may reflect the results obtained by Zeng et al. (2020), who proposed a fitness valley of intermediate size wings between two adaptive peaks represented by apterous and macropterous taxa (Stroud and Losos 2016). Complete development or complete loss of wings convey direct benefits by themselves - such as dispersal and defensive capability, increased mimicry capacity or fecundity in females (Roff 1994; Whiting et al. 2003; Zeng et al. 2020) - while brachypterous wings may be positively selected and maintained only when co-opted for non-aerodynamic purposes. For all the brachypterous species we considered, wings are associated to functions such as aposematism or stridulation, with very few exceptions such as *Hypocyrtus ornatissimus* (Brunner von Wattenwyl 1907). In the ground-dwelling Heteropterygidae, acoustic adaptations have been proposed as an initial driver of brachypterous wings reversal, with macroptery secondarily restored in species which adopted an arboreal lifestyle (Bank et al, 2021a).

Despite the known limitations of ASR in inferring precise number and position of transitions (Goldberg and Igi• 2008; Duchêne and Lanfear 2015), our analyses aim to a global evaluation of wings evolutionary patterns in the clade: reversals to brachypterous and macropterous forms were inferred in all analyses and we never recovered a macropterous root. Using the 2-state character coding scheme on the PC-Tree, extant Phasmatodea MRCA was always reconstructed as apterous even when considering different root’s prior probabilities. Using the 3-states coding on the PC-Tree recovered either a apterous or brachypterous MRCA of Phasmatodea, depending on the combination of transition rates and root assumptions, yet always rejecting a macropterous MRCA. The two ASRs where a brachypterous MRCA is found and Euphasmatodea have diversified as brachypterous-like are preferred by the AICc and both 100Sim and 100Trees SCMs point to a brachypterous root. Yet, reversals from apterous forms to brachypterous or macropterous ones are inferred only on terminal branches, an outcome without a clear biological sense.

An extant phasmid MRCA which lacked fully developed wings is consistent with paleontological evidence: extinct species representing the stem-group of all extant Phasmatodea (*e*.*g*. Susumanioidea, Archipseudophasma and Pterophasmatidae) present wings with tegmina longer than all other extinct and extant Euphasmatodea taxa (Yang et al. 2019; Yang et al. 2021; Figure 1). It is therefore possible to hypothesize that the ancestor lineage of extant phasmids presented two fully developed wing pairs and experienced either a reduction or a loss of wings. Then, subsequent reversals happened in the lineages leading to extant forms, which restored the structure in a partially different form; such differences between the derived state of a trait and its ancestral form may, in fact, represent an evidence of its reversal (Cronk 2009; Recknagel et al. 2018). Phenotypes are rarely derived from single or few genes, often the resulting from a large number of them: modifications in derived traits with respect to their ancestral form could be explained by the possible co-option of novel genes and the decay of others, along with the preservation of pleiotropic ones due to selection for other traits. In insects, wings and legs are tightly linked in a developmental perspective: their imaginal discs generated from the same group of cells and the genetic pathway that guide both structure development is largely shared (Kim et al. 1996; Cohen et al. 1993). Moreover, it has been observed that leg regeneration during phasmids development leads to smaller wings and weaker flight capability (Maginnis 2006) and that the neural structures and their functional connectivity necessary to sustain flight are conserved also in apterous forms, showing how loss of flight is not correlated to loss of associated muscles and innervations (Kutsch and Kittmann 1991). Thus, extant phasmids wings - as other reversals - could blur the boundaries between reversion and novelty, presenting a trait which is only partially built on the same genetic blueprint which produced the once lost structure.

Genomic and transcriptomic studies are contributing to elucidate the molecular consequences of trait loss and possible mechanisms associated to trait reversal (Seher et al. 2012; Carlini et al. 2013; Esfeld et al. 2018; Lammers et al. 2019) which can be - theoretically - used as informative priors to be applied in the framework of comparative methods. Standard approaches do not consider any prior assumptions on the mechanism of evolution, leveraging transitions rates estimated on the basis of trait distribution at the tips of the phylogeny and the tree itself. When there are no particular expectations on the relative probability of transitions between states, this seems a valid approach; however, an equal probability of losing and reverting back to a complex structure - *i*.*e*. made by several integrated parts - represents a strong deviation from common expectations and assumptions. While the majority of our analyses found comparable rates of losses and reversals, as can be observed from global and local approaches on the 2-states character coding strategy, we also tested the impact of different assumption on the relative probability of reversal compared to loss (Figure 5). As no study has ever explored possible mechanisms of wings reversal in phasmids, we arbitrarily tested different ratios of wings reversal to loss (from 1:10 up to 1:500, in conjunction with absolute values of the parameters consistent with their optimization in the ER and LO models. Despite a big effect of the model parameter absolute values can be observed, until a relative probability of 1:30 is assumed, reversals are consistently inferred. When reversals are assumed as more unlikely events, they are no more inferred consistently; however, even when reversals are considered highly unlikely, a scenario of irreversible evolution is never consistently supported. Previous debates on the possible reversible evolution of phasmids wings found very contrasting results, with the absolute values of the evolutionary model parameters playing a major role in defining a probability threshold above which the reversible evolution scenario is no longer supported (Whiting et al. 2003: 1:1500; Trueman et al. 2004: 1:13). Despite there is no approach to determine such a threshold (Stone and French 2003), we showed that our findings are solid to the plausible expectation of reversals being more unlikely events than losses.

## Conclusions

Altogether, the findings presented here support a dynamic and reversible evolution of phasmids wings: our analyses inferred either the absence or an extreme reduction of wings in the MRCA of extant Phasmatodea, with multiple reversals subsequent restoring the once lost structures. Neither wings or flight are found to significantly impact lineage diversification in the clade and brachyptery is recovered as an unstable state. The consistency observed with multiple complementary approaches and the scarce impact of different topological arrangements and divergence times, let us hypothesize that our findings will not be easily challenged by future phylogenetic hypotheses or comparative approaches. Nonetheless our evidences are limited to a macroevolutionary framework and complementary evo-devo, genomic and transcriptomic approaches should try elucidate the possible mechanisms underlying this phenomenon. In our opinion, rejecting Dollo’s law in this scenario is a matter of how to consider the homology of the derived and ancestral trait states. Yet, independently from the different perspectives which can be adopted on the topic, phasmids wings represent an extraordinary example of how dynamic the evolution of complex traits can be.

## Data deposition

All sequences produced have been submitted to GenBank under the accession number: MN449491-MN449962, MT077516-MT077845, MW089913 - MW089960, MW089961 - MW090026, MW138647 - MW138724, MW138492 - MW138572, MW138575 - MW138646, MW138438 - MW138491, MW528968 - MW529005, MW530366 - MW530419, MW089913 - MW089960, MW089961 - MW090026, MW138647 - MW138724, MW138492 - MW138572, MW138575 - MW138646, MW138438 - MW138491, MW528968 - MW529005, MW530366 - MW530419.

## Supplementary materials

Data available from the Dryad Digital Repository: http://dx.doi.org/10.5061/dryad.[NNNN ]

## Funding

This work was supported by Canziani Funding to AL and BM.

Acknowledgments

The authors would like to thank Mattia Ragazzini for the help in revising the character-coding tables. We also wish to thank Taiping Gao and Hongru Yang for permitting the usage of the photo of *Aclistophasma echinulatum* fossil.

## Supplementary Materials

**Supplementary Figure 1 -** The maximum clade credibility tree obtained from the the Bayesian Inference including monophyletic constrains to match the phylogenomic resolution of Tihelka et al. (2020) (PC-Tree); posterior probabilities are reported at each node. Numbers 1 to 4 represent the calibration points described in Supplementary Table 4.

**Supplementary Figure 2 -** Transition rates inferred with the 2-states (a) and 3-states (b) character coding schemes on the PC-Tree. The corresponding best-fit evolutionary models were used, respectively Equal Rates (ER) and All Rates Different (ARD). Red color highlights the transition rates which move away from brachyptery.

**Supplementary Figure 3 -** Ancestral State Reconstructions (ASRs) on the PC-Tree leveraging 2-states character coding scheme with different models of evolution and root priors.

**Supplementary Figure 4 -** Ancestral State Reconstructions (ASRs) on the PC-Tree leveraging 3-states character coding scheme with different root priors.

**Supplementary Figure 5 -** Heatmap representing the consistency of reversals inference in stochastic character mapping (SCM) analyses when different combination of model rates and relative probability of reversal are assumed. The analyses leveraged either 100 simulations on the PC-Tree (100Sim) or a single simulation on 100 randomly-sampled trees from the UC-Trees (100Trees). In (a) and (b) when consistency equals 100%, at least one reversal is inferred in each of the 100 SCM analyses for the relative model parameters combination; if consistency <100% and >0% reversals are inferred just on a fraction of the considered trees and character histories. The number of reversals and losses are reported respectively for both 100Sim (c and e) and 100Trees (d and f) analyses. NB: panel (a) is equivalent to Figure 5 in the main text.

**Supplementary Table 1 -** Species included in the analyses along with the references of the species downloaded from NCBI and the 2-states and 3-states character coding schemes. In the 2-states column, 0 represents apterous species while 1 winged ones (including both brachipterous and macropterous). In 3-states column, apterous species are coded as 0, brachipterous as 1 and macropterous as 2. In the flight column macroptery is coded as 1 while brachyptery and aptery as 0 (also including four species which are macropterous but not able to fly due to their excessive size).

**Supplementary Table 2 -** Primers sequence and PCR annealing condition.

**Supplementary Table 3 -** Maximum Likelihood model selection performed by ModelFinder with the -MERGE option and in according to the BIC score.

**Supplementary Table 4 -** Fossil calibrations used for the divergence times analysis. The calibration nodes are also reported in Supplementary Figure 1.

**Supplementary Table 5 -** AICc, -lnL, and •AICc values of the evolutionary models fitted on the PC-Tree tree with the 2-states (a) and the 3-states (b) character coding schemes. Description of the evolutionary models can be found in the main text.

**Supplementary Table 6 -** Ancestral State Reconstructions (ASRs) results for the 2-states (a) and 3-states (b) character coding schemes on the PC-Tree. Root.p = root prior probability, root.state = probability of ASRs at the root were PP = posterior probability. In (a) q01 = transition rates between apterous and winged; q10 = transition rate between winged and apterous. In (b) q01 = transition rates between apterous and brachypterous; q02 = transition rate between apterous and macropterous; q12 = transition rates between brachypterous and macropterous; q10 = transition rate between brachypterous and apterous; q20 = transition rate between macropterous and apterous; q21 = transition rate between macropterous and brachypterous.

**Supplementary Table 7 -** (a) and (b) model selection results using *SSE models on the PC-Tree, leveraging respectively the 2-states character coding scheme used for other analyses (to test for wings presence / absence) and a modified version where brachypterous species and four macropterous species which are not able to fly are coded as 0 (to test for flight capability). Details on the evolutionary models can be found in the main text.

